# Amplifying the Neural Power Spectrum

**DOI:** 10.1101/659268

**Authors:** J. Andrew Doyle, Paule-Joanne Toussaint, Alan C. Evans

## Abstract

We introduce a novel method that employs a parametric model of human electroencephalographic (EEG) brain signal power spectra to evaluate cognitive science experiments and test scientific hypotheses. We develop the Neural Power Amplifier (NPA), a data-driven approach to EEG pre-processing that can replace current filtering strategies with a principled method based on combining filters with logarithmic and Gaussian magnitude responses. Presenting the first time domain evidence to validate an increasingly popular model for neural power spectra [1], we show that filtering out the 1*/f* background signal and selecting peaks improves a time-domain decoding experiment for visual stimulus of human faces versus random noise.

## 1 Introduction

Automatic extraction / identification of electroencephalography (EEG) signal from noise is a required step towards making applications practical and less dependent on trained professionals. Standard analytic pipelines for EEG data include manual channel removal, down-sampling, artifact correction, and feature extraction.

Power spectral distributions (PSDs) are popular low-dimension summary descriptors of the high-dimensional EEG signal. In power spectra, the amplitude of a sinusoidal signal decomposition at different frequencies can be associated with underlying oscillatory properties of the time domain signal. Typical neuroscience studies predominantly compare the total PSD of standard pre-defined frequency bands (eg. *α*: 8 -12 Hz, *β*: 12.5 - 30 Hz) between groups of subjects, or for different tasks performed by a given participant. Time-frequency methods estimate power spectra computed from a narrow time-dependent sliding window to combine information from the frequency and time domains [2]. However, to simplify interpretation, the phase information that would allow returning to the time domain is discarded and information is lost. More recently, the covariance between total power and the phase of the Fourier coefficients has been used as a low-dimensional descriptor in a technique called phase-amplitude coupling (PAC) [3]. The lack of a gold standard for PAC detection has resulted in many competing methods, with only a handful providing a quantitative measure of this phenomenon[4].

Although earlier efforts to further reduce the dimensions of the power spectrum for EEG analysis suggested that no useful information could be extracted other than the moments of the PSD [5], parametric models for the PSD of EEG have been developed that decompose the spectrum into a “background” process and peaks above it that represent oscillations [6, 1, 7]. The shape of the PSD has received interest in many fields, and is commonly further simplified and expressed in terms of colours (e.g. “white noise” is often assumed, “pink noise” refers to the 1*/f* signal [8]).

Deep learning has been increasingly applied to EEG data, but best practices for pre-processing in the context of deep learning remain unclear [9]. Artifact detection or correction remains difficult and subjective, and many groups avoid pre-processing altogether by working on raw data [10], or incorporate robustness to artifacts by simulating them instead of correcting them [11]. Deep learning methods often sacrifice the interpretability of simpler linear statistical models for increased predictive power.

In this work, we introduce a data-driven pre-processing method to filter EEG recordings based on a simple parametric model of the power spectrum. Our approach combines simple feature-engineering with advanced machine learning methods and allows us to filter out the 1*/f* portion of the signal. Thus, we preserve interpretability without sacrificing performance. Replacing standard, arbitrary and manual pre-processing schemes, we design and apply digital filters based on the spectral properties of neural recordings to allow a decoding experiment in the time domain using deep learning.

More specifically, we present methods to design digital filters based on the parameters of the power spectrum, with atypical ideal transfer functions: logarithms and Gaussians. We then show how a system of filters, the Neural Power Amplifier (NPA), can be applied to filter out the 1*/f,* summarized in figure 1. We next demonstrate how the NPA allows testing the contributions of each spectral component to human neural information processing in a face vs. noise visual stimulus decoding experiment, where we compare pre-processing strategies in a deep learning investigation. Additionally, we present a method to automatically remove blinks with a strategy based on independent components of electro-oculogram (EOG) measurements.

**Figure 1:**
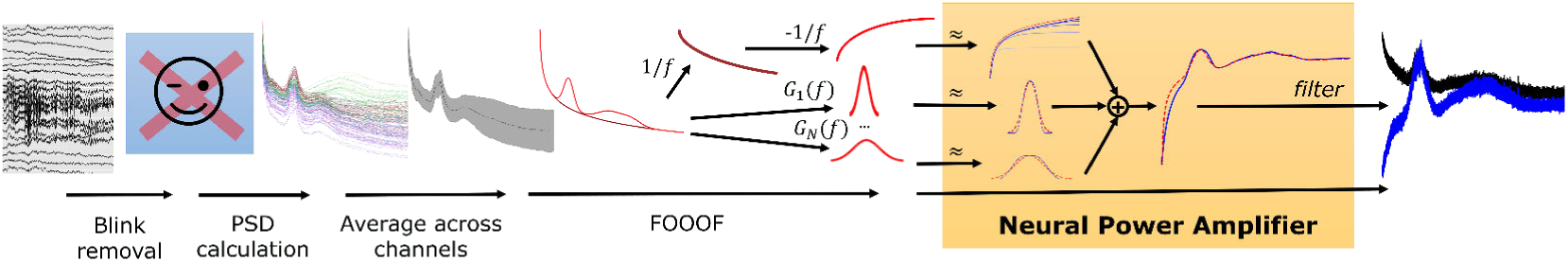
Overview of the Neural Power Amplifier method. Blinks are removed automatically, then the PSD is calculated at each electrode and averaged. The FOOOF parameterization is then approximated in digital filters by the NPA, with the logarithmic 1*/f* component normalized and negated. This system of filters is applied to the original EEG signal, which has the effect of normalizing the time series’ power spectrum.

Our cross-validated inter-subject decoding study suggests that an automatic method of blink artifact removal, in addition to NPA filtering, improves decoding accuracy in the case of face recognition vs. other visual stimulus.

## 2 Proposed Method

The main contribution of this work is the development of design methods for digital filters that approximate non-traditional ideal transfer functions, and their combination to test hypotheses based on low-dimensional descriptors without loss of information. The *χ*-*α* parametric model of the power spectrum for EEG [6] has recently been extended to include multiple peaks [1] as:

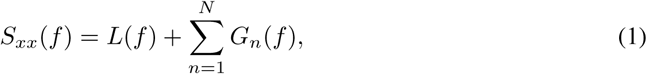

where *L*(*f)* models the 1*/f* non-oscillatory component and *n* Gaussian peaks *G*_*n*_(*f)* that often correspond to the traditional frequency bands (*α, β, δ, γ, θ*). These parametric models provide lower-dimensional descriptors for the already low-dimensional power spectrum of neural recordings and enable comparison of the parameters of the model rather than power in the traditional frequency bands. Here, we instead use these parameters to automatically design digital filters that invert the 1*/f* non-oscillatory background and select the oscillatory components so that they are not attenuated while filtering out the 1*/f*. By contrast, our method allows a return to the higher-dimensional time domain, where we can leverage the power of deep learning to test the model’s validity and estimate the effects of the parameters, as we show in section 3.2.

The application of the NPA to an EEG signal requires the following steps: (1) calculating the PSD of each electrode channel, (2) *fitting oscillations and one over frequency* (FOOOF) to a channel-averaged PSD [1], (3) designing a system of filters to approximate the logarithmic 1*/f* component of the the PSD, (4) designing a single filter to approximate each Gaussian component of the PSD, (5) filtering the EEG signal forwards and backwards with each filter to achieve zero-phase and adding their output.

### 2.1 Filter Design

Traditionally, design criteria for digital filters maximize signal transmission within a “passband” and minimize the output of signals at “stopband” frequencies. In this work, the non-ideal parts of the response in the “transition” band are essential for producing the desired logarithmic and Gaussian magnitude responses, therefore most usual design methods are not applicable. Our goal is to attenuate the 1*/f,* non-oscillatory *χ* process, while preserving other information in the multi-channel time series *x*(*t*). We require the filters to be bounded-input bounded-output stable, and to have a linear phase response. A necessary condition for stability is that the output be bounded between 0 and 1.

#### Logarithm filter

The 1*/f* process has a logarithmic spectral distribution with linear frequency, and is modeled as *L*(*f)* = *b*-*log*_10_(*k* + *f* ^*χ*^), where *b* is the offset, *χ* is the slope, and *k* is a “knee” parameter (particularly useful for modeling intra-cranial EEG) [1]. Approximating log functions is difficult with polynomials [12] and here we have the additional constraints of any approximation being realizable as a linear time-invariant digital filter. To achieve this, we negate *L*(*f)* and normalize this infinite-range function by dividing it by the maximum in the frequency range where the filter is to be applied, which occurs at the Nyquist frequency (*f*_*nyq*_) and clipping negative values to 0:

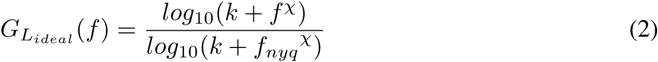

We approximate 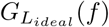 by applying a system of *M* order-1 high-pass Chebyshev filters in parallel and adding their output. Each Chebyshev filter *m* has a magnitude response

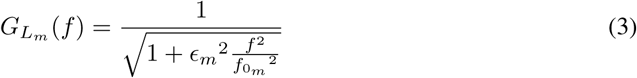

and approximates the value at half the maximum of the remaining ideal logarithm transfer function. To achieve this, each filter *m* ∈*M* is designed with a critical frequency 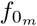, selected where the monotonically increasing 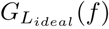 reaches half of the remainder of the gain not approximated by the previous stages,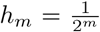. For *m* = 1, since 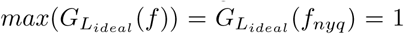, the first stage approximates up to 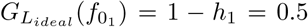. The critical frequency for filter *m* is given 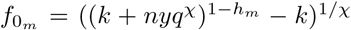. For the first stage, this corresponds to the -3*dB* cutoff reminiscent of traditional filter design methods. The ϵ_*m*_ parameter traditionally controls the maximum allowable ripple in the pass-band. For single-tap filters, there is only a single “ripple” below the maximum gain of 1, so it corresponds to a tuning parameter for the slope of the transition band. We choose 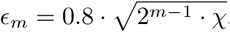, which produces a transfer function based only on *χ* and *k*, properties of the EEG signal:

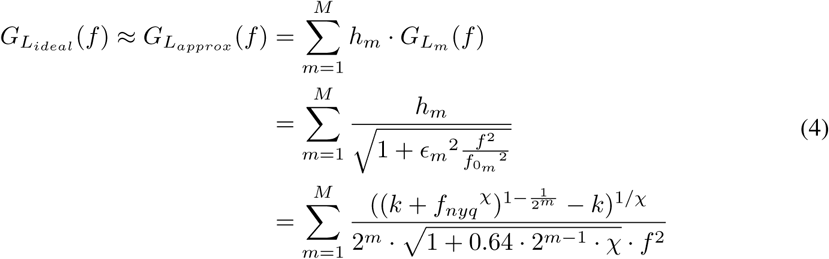

#### Gaussian filter

The oscillatory components of neural recordings are thought to produce peaks in the power spectrum, the most prominent of which are typically the “alpha” waves between 8-12 Hz. These peaks are modeled in a single power spectrum by a variable number *N* of Gaussian functions, where each peak *G*_*n*_ is described by an amplitude *a*_*n*_, mean *µ*_*n*_ and standard deviation *σ*_*n*_. As such, the ideal response of a filter that selects oscillatory components of the EEG signal *x*(*t*) is given by 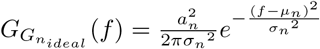. This ideal magnitude is a *squared* Gaussian because, as with the logarithm, this filter will be applied forward and backwards to achieve zero phase.

In contrast to the logarithmic case, where we used many single-tap filters for the approximation, here we use a single filter with many taps to obtain the desired magnitude response. The Remez exchange algorithm [13] is used to produce coefficients *b* for optimal filters with approximately Gaussian magnitude responses 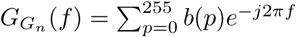. Our approach requires the specification of two stopbands, from *f* ∈ [0, *µ*_*n*_ - 3*σ*_*n*_] and *f* ∈ [*µ*_*n*_ + 3*σ*_*n*_, *f*_*nyq*_], and a single passband. The edges of the passband are chosen to be at -3*dB* of the maximum amplitude, which occurs at *µ*_*n*_, so the passband edges are at 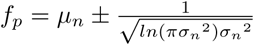

Thus, the total ideal magnitude response of the Neural Power Amplifier (NPA) consists of the negated logarithmic 1*/f* process and a filter for each Gaussian present in *x*(*t*):

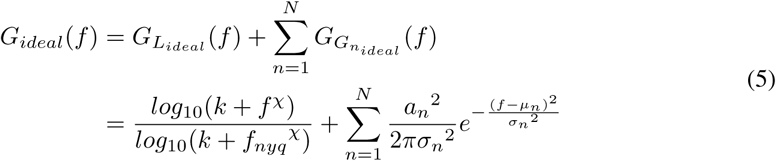

This is approximated here by equation 6, where all terms are estimated in a data driven manner from a simple parametric model of a neural power spectrum:

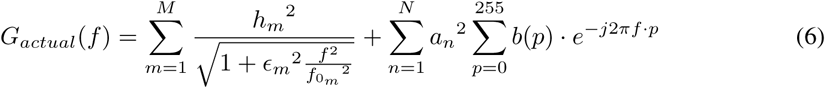

### 2.2 Data and Pre-Processing

Application of the NPA requires parameters from power spectra of EEG signals. Typically, EEG data are heavily pre-preprocessed with aggressive filtering, down-sampling and manual channel/artifact rejection [14]. This pre-processing is highly time-consuming and subjective based on what follow-up analysis is being done, and even what programming language is used (!). Such an approach is not reproducible, or even feasible for large volumes of data.

#### Data

The data used in this work were previously collected to test reproducibility of Event Related Potentials^1^ (ERPs) of a visual face stimulus, where 4 participants were each scanned in 10 separate 1-hour sessions. Images of human faces or random noise were presented for 82 ms each, and EEG was recorded at 512 Hz using a BioSemi system with 128 EEG electrode channels and 4 EOG electrodes.[15]

#### Line noise artifacts

Power line noise (50 Hz and harmonics here) was attenuated using the *cleanline* plug-in [16] for the MATLAB-based EEGLAB [17]. Although *cleanline* was not found to completely remove spectral peaks, we prefer this method to notch filtering, which might remove signals of interest that overlap in the *γ* band (30+ Hz). All other analyses were done using Python 3.7, Scipy 1.2 and MNE 0.17.1 [18, 19].

#### Eye blink artifacts

Eye blinks are the strongest stable artifact detectable in EEG. Approximately 39 seconds (20,000 samples) of EEG were selected and bandpass filtered between 1 and 40 Hz, then downsampled to 128 Hz. The filtered timeseries subset was decomposed with principal component analysis (PCA), and the components that explained up to 98% of the variance were further decomposed using independent component analysis (ICA). The independent component that explained the most variance of the right vertical EOG channel (channel EXG1) was assumed to correspond to an eye blink, and that component was subtracted from the raw (unfiltered, original sampling rate) signal.

An example of the spatial distribution of a blink IC, its correlation with the EOG channel, and its contribution to the time series magnitude is shown in figure 2.

**Figure 2:**
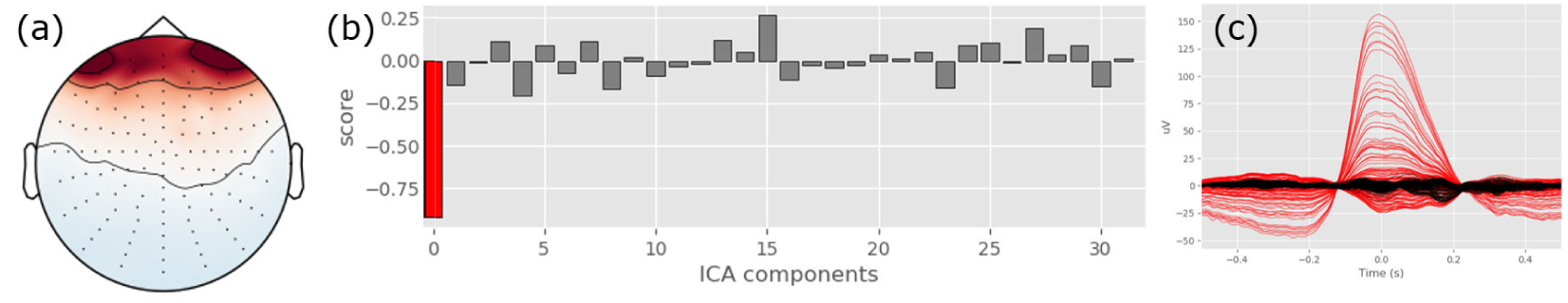
Blink detection and removal from a single representative session. Spatial distribution of signal removed shown in (a), correlation of each independent component with EOG shown in (b), with the component having highest correlation shown in red. The average signal before (red) and after (black) ICA correction is shown in (c)

#### Power Spectra

PSDs were computed over each 1-hour session for each electrode using a multi-taper method with 7 tapers [20]. The channel-averaged spectrum was then parameterized using FOOOF [1], with a minimum peak amplitude of 0.05 and peak width constrained to *σ*^2^ ∈[3Hz, 15Hz], including fitting the background “knee” parameter *k*, for the portion of the power spectrum between 0.1 and 45 Hz.

## 3 Experiments and Results

The experiments in this work were performed to verify that (1) the Gaussian and logarithmic filters provided good approximations to the ideal magnitude responses, and (2), that filtering out the 1*/f* process allows easier single-trial decoding of visual stimulus.

### 3.1 Neural Power Amplifier Filters

The coefficient of determination (*r*^2^) was used to evaluate how well the magnitude response for our filters fit their assumed corresponding ideal transfer functions. The logarithmic filter approximation provided a very good fit to an ideal logarithm, as shown in figure 3. We found that improvements to the *r*^2^ gains above *M* = 5 were negligible.

**Figure 3:**
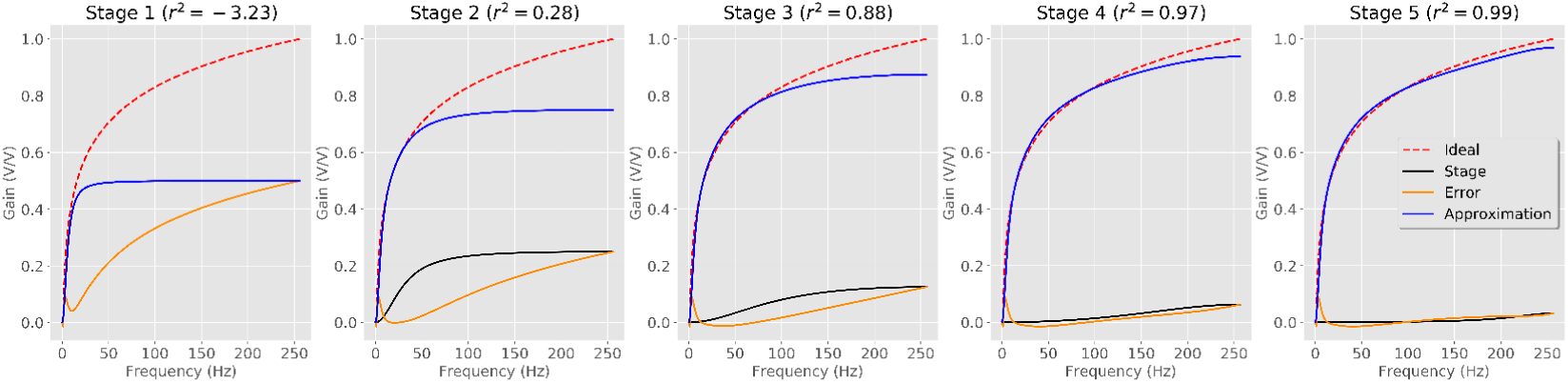
Logarithmic transfer function approximation. The first plot shows a single level of approximation, and each successive plot adds another filter to the total approximation.

The logarithmic component contributes more to the total *r*^2^ than the Gaussian components, as it has significant power that spans the entire power spectrum, while the Gaussian peaks occupy less bandwidth. This can be seen in figure 4, where the logarithmic filter approximation spans up to *f*_*nyq*_, while the Gaussians found by FOOOF extend just past 45 Hz. Figure 4 also shows good qualitative fits to the ideal Gaussians, which begin to worsen as their standard deviation spread out.

**Figure 4:**
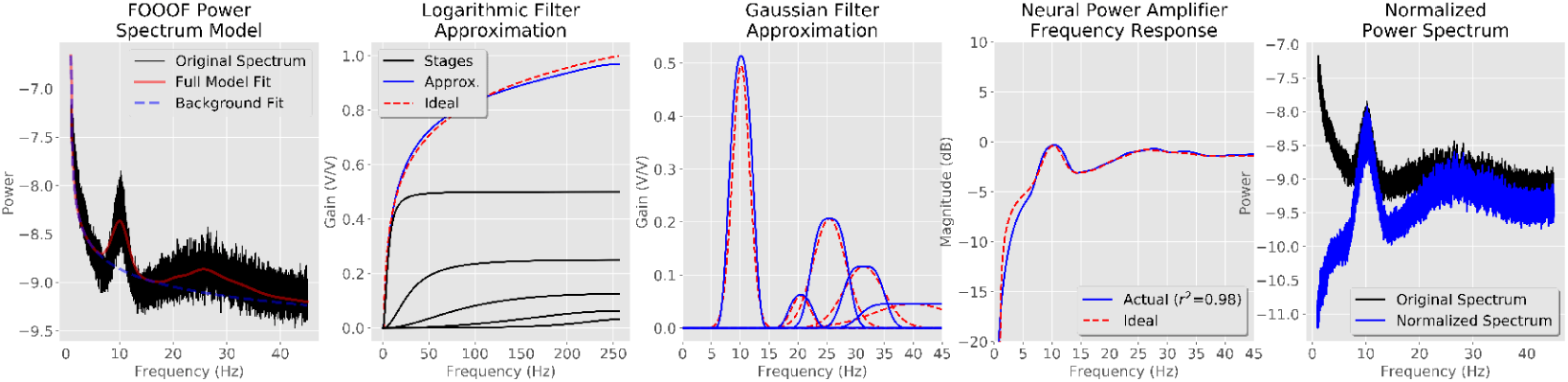
Neural Power Amplifier filter response results. The first panel shows the FOOOF model fit. The second panel shows the NPA approximation for the logarithm based on the FOOOF model. The third panel shows the magnitude response (blue) for all of the Gaussian peaks found by FOOOF (red). The fourth panel shows the total magnitude response in decibels. The fifth panel shows the power spectrum of the original time series (black) and the NPA amplified time series (blue), demonstrating that the 1*/f* has been effectively removed while amplifying the peaks.

The FOOOF model usually provided a good model fit for each EEG recording session’s PSD, and the NPA, based on the FOOOF *χ* and *k* parameters, produced reasonable approximations as well, with *r*^2^ results for each participant shown in figure 5. We observed that for some sessions, the FOOOF model determined a negative value of *k*. This presented an unexpected edge case for the Neural Power Amplifier, as digital filters are undefined at negative frequencies. However, in our data, the negative values of *k* were small enough (*<* 1 Hz in all sessions) that the *k* = 0 approximation still produced reasonable NPA model fits.

**Figure 5:**
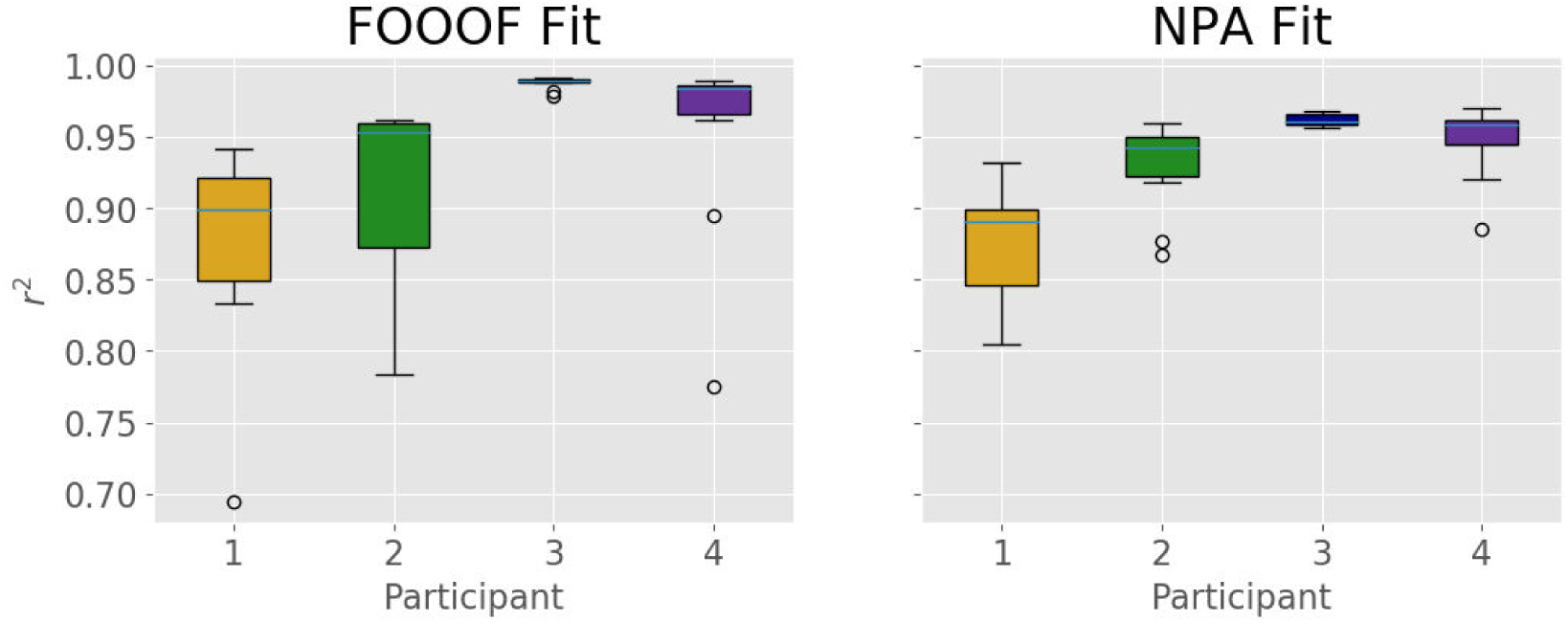
Resulting coefficient of determination for the FOOOF parametric model of power spectra for each participant (left) and magnitude response function of the Neural Power Amplifier (right).

### 3.2 Neural Decoding

Our final evaluation of the proposed Neural Power Amplifier compares it against more standard pre-processing. Ground truth in human neuroimaging separating brain signals and noise does not exist, so we conduct a neural decoding experiment for known sensory stimuli and use decoding accuracy as a heuristic to measure the appropriateness of the method. We compare these conditions to test the validity of the proposed Neural Power Amplifier method as a way to remove inter-subject variability for easier decoding of neural stimulus. As such, we can also test the parametric model on which the NPA is based to provide some evidence for whether these parameters are meaningful. Finally, it allows a novel method to investigate the true nature of the 1*/f* process and determine how it contributes to neural computation, a mechanism which remains poorly understood.

We compare several leave-one-subject-out supervised machine learning experiments, using a 16-filter Long Short-Term Memory (LSTM) neural network with a max-norm constraint of 2 on the recurrence to predict whether subjects are looking at images of a human face, or viewing random noise patterns [21]. The input to the LSTM consists of epochs from the 128 EEG electrodes described in section 2, which corresponded to 667 time points per trial. Mini-batches consisted of 1024 samples, and the model was trained for 20 epochs, sampling 3 mini-batches per epoch from the training set. The network is trained using stochastic gradient descent with a learning rate of 0.0002, and was initialized at the same random seed for four pre-processing conditions on *x*(*t*): (1) the proposed Neural Power Amplifier method, (2) band-pass filtering from 1-40 Hz, (3) high-pass filtering at 1 Hz, and (4) raw signal. For all types of pre-processing except the raw signal, we apply our automatic ICA-based automatic blink artifact removal. Decoding results are shown in figure 6 over the course of training for each fold, as well as the test decoding accuracy.

**Figure 6:**
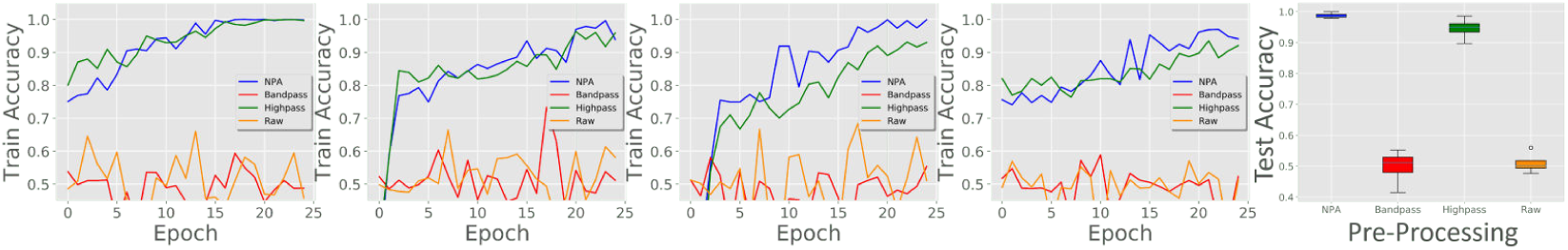
Results from human faces vs. random noise visual stimulus decoding experiment. Four pre-processing methods are compared: the Neural Power Amplifier (blue), bandpass filtering (red), highpass filtering (green), and raw EEG (orange). Plots of accuracy are shown for each fold of training (left) and final testing accuracy (right).

The Neural Power Amplifier outperformed all other methods, achieving over 98% test accuracy. While the very common bandpass filtered EEG could not be decoded, the highpass filter achieved good results as well, suggesting that high frequency information beyond what is normally considered is indeed important in the context of visual human face processing.

Code to reproduce all experiments in this work can be found at https://github.com/crocodoyle/good-noise.

### 3.3 Event Related Potentials

EEG experiments are often evaluated using ERPs, which average the EEG signal thought to be produced by a particular stimulus. Many trials are often required to observe stable spatiotemporal patterns across the brain. Human faces produce some of the most reliable ERPs, with the N170 signal producing a negative voltage deflection at 140 - 200 ms in the occipito-temporal electrode sites, and positive voltage in fronto-central regions following human face stimulus presentation[22].

For comparison with traditional methodologies, we show the difference between faces and noise stimuli in a single participant’s ERPs in figure 7. The average voltage recorded across all trials is normalized by the variance across trials for each of the face and noise stimuli. The normalized noise stimulus is subtracted from the normalized face stimulus ERP to produce a visualization of the most stable pattern differences between the different pre-processing conditions under test.

**Figure 7:**
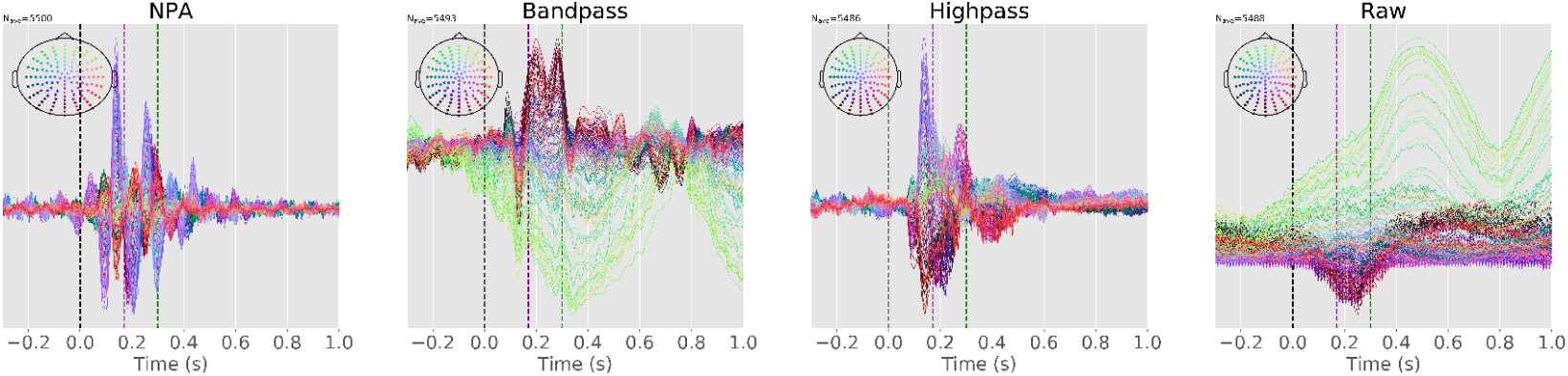
Normalized differences in ERPs for face presentation stimuli for the four pre-processing conditions. Stimulus onset (black), N170 (purple) and P300 (green) times are shown as dashed lines.

While it is not surprising that the NPA produces relatively stable *α* waves (we explicitly amplified these), it is interesting that this pattern should appear in a normalized difference between two conditions. In these ERP differences, the N170 negative deflection is identifiable in the purple electrodes’ response in the NPA, while the P300 is more obvious in the green of the raw signal. Furthermore, the N170 appears to be synchronized with the *α* band in the NPA response, while the P300 is not. These qualitative observations are consistent across the four subjects, and suggest that the NPA method has the potential to further investigate the underlying processes that generate these phenomena.

## 4 Conclusion

With this work, we introduce a novel, data-driven, automatic alternative pre-processing strategy to the arbitrary pre-processing approaches in common use for studying EEG. We employed this pre- processing approach to add quantitative evidence for the validity of the *χα* parametric decomposition of the power spectrum, and expand methods of neuroscience inquiry by allowing a return to the time domain based on spectral signal characteristics.

We introduce a new tool to investigate a phenomenon long observed, but poorly understood. The non-oscillatory 1*/f* process is certainly not purely noise, but we find with this work that in the context of fast neural processes like facial recognition, the 1*/f* might contain less information than the oscillatory bands. Additionally, our approach shows preliminary evidence that the long-observed N170 response is related to *α* waves, while the P300 is not (figure 7).

There are several limitations of our implementation of the NPA method presented in this work. The Remez-based algorithm for the Gaussian-response filter fails to converge for Gaussians with larger standard deviations, where the transition band of the resulting filter becomes too wide, and a cruder approximation is necessary. Also, FOOOF’s optional knee parameter cannot be well-modeled using our system of single tap filters approach for large *k*, as the magnitude response increases monotonically from 0.

Our approach averages the PSD over all EEG channels, but a per-channel time-varying filtering strategy might be more relevant. In this initial study, we were unable to find a real-time filtering strategy that satisfied our other design parameters. Filtering forwards and backwards solves the problem of phase distortion being introduced by infinite impulse response filters, but also requires that the entire signal of interest be acquired before filtering can begin in the backwards direction. While a windowing strategy might enable near real-time application, the delay introduced by filtering requires a minimum number of samples before all frequency components are present at the output. However, the Gaussian approximate filter (that theoretically does not need to be applied backwards because of the linear phase response of Finite Impulse Response filters) still requires 256 samples, which amounts to 0.5 seconds in our data.

Calculation of the PSD is the most computationally expensive operation required by the Neural Power Amplifier method. To obtain the highest precision, we compute PSDs for each EEG channel using the most accurate (and time-consuming) method over an entire hour of signal. This is not feasible for real-time applications, which would require the selection of a relatively wide window size in order to have enough resolution to represent the low-frequency components necessary for the PSD parametric model estimation. While simpler, less expensive methods for PSD estimation exist such as Welch’s method [23] or periodograms, it is unclear whether they produce good enough estimates to enable good FOOOF model fits.

Despite its limitations, the Neural Power Amplifier is a novel tool with broad applicability that allows the combination of previous findings in both frequency and time domains, as well as previous work in neuroscience (neuronal decoding), computer science (deep learning) and electrical engineering (signal processing). We combine linear, interpretable filtering of canonical frequency bands with state of the art deep learning techniques in an automatic, reproducible manner. Our approach is easily adaptable not only to other EEG experiments, but also to other neuroimaging modalities like magnetoencephalography, or even functional magnetic resonance imaging, which have much lower frequency but also exhibit the 1*/f* frequency behaviour. While the nature of this mysterious signal remains unclear, we hope that this tool will provide investigators a new avenue to uncover its role in information processing.

## 5 Acknowledgements

We thank Dr. Guillaume Rousselet, Hanna Isotalus and Sean Henderson especially for collecting, documenting, and sharing data, without which this research would not have been possible. We thank Dr. Pedro Valdés-Sosa, Dr. Sylvain Baillet and Dr. John Griffiths for the valuable discussions they provided, as well as Dr. Pamela K. Douglas for inspiring this line of research. This work was supported by the Fonds de Recherche du Québec - Santé, the Canada-China-Cuba Axis, and the NVIDIA Academic Hardware Grant.

## Footnotes

1 ERPs are averages of neural recordings over many trials, sessions, or participants, and represent stable patterns that the stimulus is thought to reliably produce.

